# Beyond Gatekeeping: Efflux Pumps Remotely Destabilize Cytoplasmic Drug–Target Interactions by Limiting Rebinding

**DOI:** 10.1101/2025.08.06.668934

**Authors:** Subrata Dev, Keiran Stevenson, Dai Le, Minsu Kim

## Abstract

Bacterial efflux pumps are major contributors to multidrug resistance, classically described as “gatekeepers” that reduce drug entry. Here, we uncover an equally potent post-entry mechanism of efflux pumps, revealing that their function extends deep into intracellular drug-target interactions. Using quantitative live-cell fluorescence imaging, we monitored the activity of major efflux systems in *Escherichia coli* and *Pseudomonas aeruginosa* with Hoechst (HCT), a DNA-binding inhibitor. As expected, more HCT entered efflux-deficient cells than efflux-active cells. Unexpectedly, the intracellular HCT in efflux-deficient cells also bound DNA more stably, resulting in higher binding affinity. Statistical-physics-based modeling and experimental testing show that that unlike in dilute in vitro conditions, drug molecules that unbind from their targets in intracellular environments rapidly undergo successive rebinding, prolonging the lifetime of the drug-target complex. However, efflux pumps counteract this effect by suppressing rebinding, thereby kinetically destabilizing drug-target interactions and lowering the binding affinity. This kinetic destabilization acts synergistically with canonical gatekeeping to broaden and amplify drug resistance. Our findings reveal an underappreciated biophysical mechanism of efflux pumps, expanding the classical biochemical view of how efflux pumps confer multidrug resistance.

## Introduction

Antibiotic resistance is an increasingly serious threat to public health, as existing antibiotics lose their potency and become clinically ineffective ^1^. This challenge is further compounded by the slow pace of new drug development. Most antibiotics act by binding to intracellular macromolecules, thereby disrupting cellular functions. Resistance can arise through genetic mutations that alter the structure of either the drug or its target, reducing binding affinity and diminishing antibiotic efficacy ^2^.

Recently, efflux pumps have been recognized as major contributors to multidrug resistance ^3^. Ubiquitous across bacterial species, these transporters underlie a broad spectrum of innate resistance to antibiotics ^4,5^. Additionally, their activities provide a temporal window for bacteria to mutate and acquire additional resistance genetically ^6^. As such, understanding the mechanistic basis of efflux pump function is critical for developing new strategies to counter antibiotic resistance.

It is widely accepted that efflux pumps act as permeation barriers ^4^. Embedded in the cell membrane, they capture incoming drug molecules and expel them into extracellular space. Efflux pumps are classified into five superfamilies, based on their molecular characteristics ^7^. Among them, the resistance-nodulation-division (RND) family plays a particularly important role in drug resistance due to its broad substrate specificity, extruding a wide range of structurally diverse drugs ^8^. The upregulation of RND pumps is common among clinical isolates and is strongly associated with multi-drug resistance ^9,10^.

The AcrAB-TolC system in *Escherichia coli* has served as a primary model for studying RND efflux pumps ^11^. Typical of RND pumps, it forms a tripartite complex composed of the outer-membrane channel (TolC), the inner membrane transporter (AcrB), and the periplasmic adaptor protein (AcrA), thereby spanning the entire cell envelope. The AcrAB-TolC system is widespread in Gram-negative species, including many pathogens ^12^. Given its central role in drug resistance, several inhibitors are under investigation to block its activity and restore the efficacy of existing antibiotics ^13–15^.

However, efflux pumps do not completely prevent drug molecules from reaching their intracellular targets. As external drug concentrations increase, more molecules bypass the membrane-bound pumps and accumulate in the cytoplasm, where they interact with intended targets.

In the past, drug-target interactions were extensively studied to predict the cytotoxic effects of drugs. Binding affinity is computationally modeled based on chemical structure and thermodynamics and tested in vitro using purified targets ^16,17^. These measurements have guided medicinal chemistry efforts to optimize drug molecules for higher affinity. In recent years, the mean lifetime of drug-target complex, quantified as the inverse of the unbinding rate *k*off, emerged as an additional key determinant of cytotoxicity ^18–20^. The complex lifetime matters because a drug molecule is active only when it is bound to its target. In the past, however, drug-target interactions were mostly studied in vitro. These studies do not fully capture the spatial and temporal dynamics of interactions within the native environment, i.e., within cells ^21^.

In this study, we investigated drug-target interactions in live bacterial cells, with a particular focus on the role of efflux pumps. To probe these dynamics, we used benzimidazole, one of the most common structural motifs in FDA-approved drugs ^22,23^. Due to its strong affinity for nucleotide bases ^24–29^, it is being widely adopted in medicinal chemistry to develop anti-microbials, anti-viral, anti-cancer, and anti-inflammatory agents ^30,31^.

Among these compounds, a benzimidazole derivative known as Hoechst 33342 (HCT) ^32^ exhibits physicochemical properties of drug molecules. It has a molecular weight (562 Da) and lipophilicity (LogP ≈ 2.7, based on SwissADME ^33^) typical of orally active small-molecule drugs ^34–36^. It is weakly basic at physiological pH ^37^. These properties allow HCT to cross the lipid membranes and accumulate in the cytoplasm, where it binds specifically to DNA and inhibits replication and gene expression ^38^. Notably, many of the same chemical properties are also recognized by efflux systems ^39,40^. Therefore, HCT has been widely used as a functional reporter of efflux pump activity across different bacterial species ^41–43^.

HCT’s fluorescence provides a robust method for studying drug–target interactions quantitatively ^44–47^. It fluoresces upon binding to DNA and loses fluorescence upon unbinding. We previously showed that spatial HCT intensity profiles match the spatial intensity profile of DNA-binding fluorescent proteins, demonstrating that HCT fluorescence inside the cell results primarily from its binding to DNA ^48^, in agreement with earlier biochemical studies ^32^. Therefore, HCT–DNA interactions provide a useful and tractable model for investigating drug–target binding and unbinding dynamics inside live bacterial cells.

In this study, we leveraged these properties of HCT and expanded the finding using physics-based modeling to analyze drug–target interactions inside intact cells, which shows that the effects of efflux pumps go beyond gatekeeping. First, we quantified the intracellular HCT-DNA bound state by measuring HCT fluorescence intensity in *E. coli*. We exposed efflux-deficient (Δ*tolC*) cells to a different external HCT concentration than that used for wild-type (WT) cells to equalize the intracellular level of HCT-DNA complex across strains. The resulting concentration difference could not be explained by the gatekeeping role of efflux pumps alone, instead pointing to different intracellular drug–target kinetic and/or equilibrium constants between the efflux-active and deficient strains. Indeed, our additional experiments confirmed that the efflux pumps significantly alter the apparent binding affinity, *K*D, by modulating the unbinding rate *k*off. This was surprising, given that efflux pumps are localized to the membrane and physically distant from HCT’s cytoplasmic target, DNA. To explain how efflux pumps remotely act to modulate cytoplasmic drug-target interactions, we used a statistical mechanics model of partially reflected Brownian motion, which reveals a kinetic competition between intracellular diffusion and outward permeation of drug molecules. Computational simulations showed that drug molecules, upon dissociating from their targets, often rebind. Efflux pumps facilitate their export and reduce the likelihood of rebinding, thereby altering the apparent drug–target affinity and unbinding rate without affecting the drug-target thermodynamic stability. We validated this prediction experimentally by measuring HCT–DNA rebinding in both *E. coli* and *Pseudomonas aeruginosa*, another species known for its high efflux activity. This previously unrecognized long-range kinetic mechanism reduces drug-target engagement to a degree comparable to canonical short-range gatekeeping. Only by accounting for the synergy between both mechanisms could we fully explain the reduced HCT–DNA occupancy in WT. Analyzing the diffusion constants and membrane permeability of small-molecule drugs in our model suggests that this long-range kinetic mechanism is applicable to a wide range of drug molecules. Coupled with the broad substrate specificity of efflux pumps—particularly those of the RND family ^8^— this finding provides new physical insights into how efflux pumps function to restrict drugs’ access to their intracellular targets and drive multi-drug resistance.

## Results

### Efflux pumps reduce intracellular HCT accumulation

We exposed wild-type (WT) *E. coli* to a range of external HCT concentrations and imaged the cells using fluorescence microscopy. Increasing external HCT concentrations led to higher intracellular fluorescence, with the relationship remaining linear at lower doses (Supplementary Fig. 1). However, beyond 10 µM, further increases in external HCT resulted in minimal changes in intracellular fluorescence, indicating that its target (DNA) is nearly saturated.

To avoid this saturation regime, we used 1 µM HCT for WT cells in subsequent experiments. To characterize the temporal accumulation of HCT on its target (DNA), we measured fluorescence intensity *F*(t) at various timepoints after HCT addition (time zero). As shown in Fig. 1a, *F*(t) increased initially over time and reached a steady-state plateau, denoted as *Fs*.

**Figure 1.**
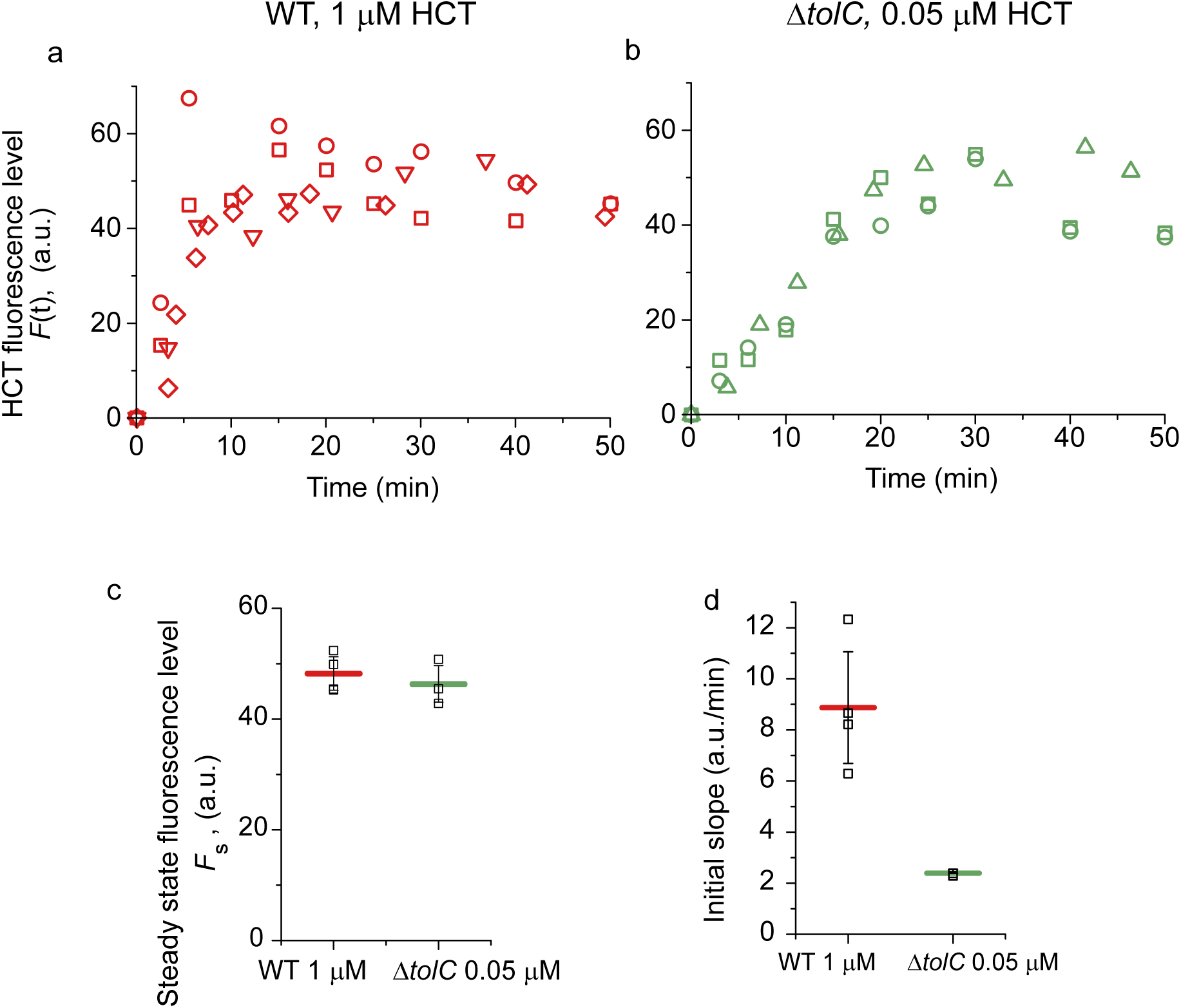
Intracellular HCT fluorescence intensity in live *E. coli* cells. a,b) HCT was added to *E. coli* cultures at time zero. The intracellular fluorescence intensity, *F*(t), initially increased and reached a steady-state level. 0.05 µM of HCT was added to a culture of Δ*tolC* cells to match *Fs* for WT at 1 µM. Three independent replicates were performed; different symbols represent different replicates. At least 30 cells were analyzed in each replicate. c) Data points between 20 and 50 minutes were averaged to obtain the steady-state fluorescence level, *F*s. Open squares represent individual values from independent replicates. A two-tailed *t*-test (with two-sample unequal variance) comparing *F*s between WT and Δ*tolC* strains yielded *p* = 0.556, indicating no significant difference. d) Data points between 0–6 minutes for WT and 0–15 minutes for Δ*tolC* were fit with linear regression to calculate the initial rate of increase. A two-tailed *t*-test (with two-sample unequal variance) comparing the slopes between WT and Δ*tolC* yielded *p* = 0.014, indicating a significant difference in the slope.

To test whether HCT is a substrate of the AcrAB-TolC efflux pump, we analyzed strains lacking *acrA*, *acrB*, or *tolC*. All knockout strains showed significantly elevated *Fs* compared to WT (Supplementary Fig. 2), indicating that HCT is a substrate for the AcrAB-TolC system, consistent with previous studies ^41–43^. Deletion of any single component—*acrA*, *acrB*, or *tolC*— was sufficient to increase *Fs*. Based on this, we used the Δ*tolC* strain for all subsequent experiments.

### Reduction in drug permeation by efflux pumps alone is not sufficient to explain their effects on intracellular HCT levels

When studying the efflux activity, it is typical to compare the responses of efflux-active and deficient strains to the same drug concentration. However, at 1 µM HCT, the steady-state fluorescence intensity (*Fs*) in the Δ*tolC* strain reached a level close to the target saturation (Supplementary Fig. 2)—a regime we deliberately avoided in the WT (Supplementary Fig. 1). This implies that, at the same external concentration, the two strains experience distinct intracellular drug regimes, complicating direct comparison.

This discrepancy introduces additional complications. Given the inhibitory effects of HCT on gene expression and DNA replication ^38^, increased HCT binding is expected to slow cell growth. Indeed, we observed a significant reduction in the growth rate of the Δ*tolC* strain at 1 µM HCT, whereas WT growth remained unaffected (Supplementary Fig. 3). Differences in growth rate complicate quantitative comparisons of intracellular fluorescence between WT and Δ*tolC*. For example, different growth rates mean different rates of cell volume increase, hence different dilution rates of intracellular drug molecules, which can alter steady-state intracellular drug concentrations ^49^. The alteration in the intracellular drug concentration can in turn affect the growth rate, forming a feedback loop ^48,50–52^, that adds complexity to the interpretation of steady-state fluorescence measurements.

To avoid these confounding effects, we instead adjusted the external HCT concentration in Δ*tolC* to match the intracellular fluorescence levels observed in WT. Through titration (Supplementary Fig. 4), we found that 0.05 µM HCT in Δ*tolC* yielded an *Fs* value closely matching that of WT cells treated with 1 µM HCT (Fig. 1ab), with no statistically significant difference (Fig. 1c caption). At this lower dose, Δ*tolC* growth was unaffected (Supplementary Fig. 3). This large reduction in the external concentrations (20-fold reduction, from 1 µM to 0.05 µM) required to achieve the same number of DNA-bound HCT molecules underscores a significant role of the efflux pumps in drug resistance.

A natural hypothesis to explain this difference in the external concentrations is that the AcrAB– TolC system effectively reduces HCT entry into the cell by a factor of 20. This idea can be formalized mathematically (Supplementary Text, section 1). Briefly, let *P*inward denote the effective permeability governing drug influx. The total influx of drug molecules is given by *P*inward×*C*out. Deletion of *tolC* will allow more drug molecules to enter the cell, effectively increasing *P*inward. If this permeability increases by 20-fold in Δ*tolC*, then with a reduction in the external concentration *C*out by the same factor, the total influx rate, *P*inward×*C*out, would remain unchanged.

Therefore, with this gatekeeping mechanism alone, once the external concentration decreased by 20-fold, intracellular HCT accumulation dynamics should be the same between WT and Δ*tolC*. However, we found that even though both strains eventually converge to the same steady-state value, their initial accumulation slopes differed significantly (Fig. 1a-c).

We next analyzed the slopes to deduce a difference in effective *P*inward between WT and Δ*tolC*. We assumed that once drug molecules enter the cells, their biochemical interactions with the intracellular targets are identical in WT and Δ*tolC* strains. With this assumption, the difference in the initial slope in drug accumulation is primarily governed by the total influx rate, *P*inward×*C*out, as discussed above. As shown in Fig. 1d, the slope in WT is higher by 3.7-fold than in Δ*tolC*. Given that the external concentration differed by 20-fold, this implies that the efflux pumps effectively reduce *P*inward by 5.4-folds (=20/3.7, see Eq. S12 in Supplementary Text).

### The HCT-DNA complex is more stable in Δ*tolC* than in WT

This 5.4-fold reduction in *P*inward by efflux pumps underscores their gatekeeping role, but also raises a question: Why was a 20-fold difference in external HCT concentration required to equalize intracellular levels between WT and Δ*tolC*Further analysis of the accumulation kinetics, *F*(t), offers a clue. Earlier, we assumed that once drug molecules enter the cell, their interaction with intracellular targets should be identical in WT and Δ*tolC* strains. However, this assumption is challenged by the observation that the Δ*tolC* strain reached the steady state substantially later than WT (Fig. 1a–b). Specifically, according to standard drug–target kinetics (Supplementary Text, section 2), the time required to reach a steady state is primarily governed by the unbinding rate, *k*off, which quantifies how quickly the drug–target complex dissociates ^16,19,20^. Based on this argument, the slower approach to steady state in Δ*tolC* suggests a lower *k*off, i.e., a more stable HCT–DNA complex.

To test this hypothesis, we directly measured *k*off using a washout experiment. We first allowed intracellular fluorescence to reach a steady-state level. As described above, WT and Δ*tolC* cells were exposed to external HCT concentrations differing by 20-fold, such that both strains exhibit a similar number of DNA-bound HCT molecules. Cells were then washed and transferred to HCT-free media, and the decline in intracellular fluorescence was monitored over time (Fig. 2a). This decline followed first-order kinetics (linear on a semi-log scale). The decline was significantly slower in the Δ*tolC* strain (Fig. 2a). We determined *k*off from the slope of this decline, which shows that it is ∼4-fold lower in Δ*tolC* relative to WT (Fig. 2b), consistent with our prediction.

**Figure 2.**
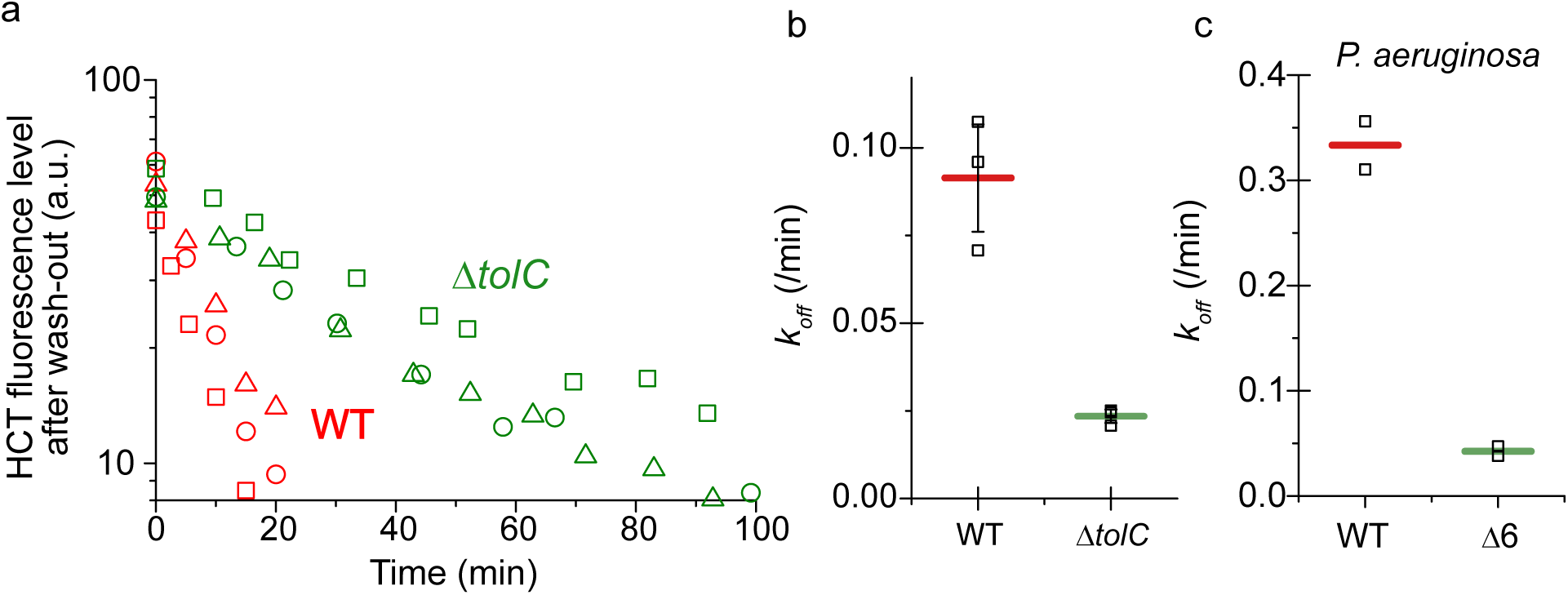
Decrease in intracellular HCT fluorescence intensity after HCT wash-out. (a) After allowing HCT fluorescence to reach its steady-state level, we washed *E. coli* cells and suspended them in HCT-free media. At least 30 cells were analyzed in each experiment. Three biological replicates were conducted (different symbols). Slower decline in fluorescence in Δ*tolC* cells than in WT indicates slower dissociation of the HCT–DNA complex. (b) The slope of this decline was analyzed to quantify *k*off. Open squares represent individual values from independent replicates. Horizontal bars and error bars indicate the means and standard deviation from the replicates. *k*off was ∼4-fold lower in Δ*tolC* compared to WT. (c). The same trend was observed in *Pseudomonas aeruginosa*: strains deficient in efflux pumps (PAO1Δ6 ^54^: Δ*mexAB-oprM,* Δ*mexCD-oprJ* Δ*mexEF-oprN* Δ*mexJKL,* Δ*mexXY,* Δ*triABC*) exhibited significantly lower *k*off values than WT (PAO1).

This difference in unbinding rates can be interpreted in terms of drug–target complex lifetime, defined as the average time a drug molecule remains bound to its target and given by the inverse of *k*off 16,53. In WT cells, the HCT–DNA complex lifetime was 10.9 minutes, whereas in Δ*tolC*, it extended to 42.6 minutes. Therefore, the complex is more stable in the efflux-deficient strain.

We tested the robustness of our findings through several additional experiments. In our initial measurements, samples were taken at multiple time points from the same culture, meaning different cells were imaged at each point. To confirm that the observed differences in *k*off were not due to population variability, we used time-lapse microscopy to monitor fluorescence decay in individual cells over time. These single-cell measurements yielded *k*off values consistent with the population-level data for both strains (Supplementary Fig. 5), demonstrating that the unbinding kinetics are representative at the single-cell level.

Next, we assessed whether *k*off depends on external HCT concentration. Across a range of doses, we found only minor variation in *k*off, and the Δ*tolC* strain consistently exhibited a significantly lower unbinding rate than WT (Supplementary Fig. 6), indicating that this effect is largely independent of external drug concentration.

Finally, we tested whether this efflux-dependent modulation of *k*off also occurs in other bacterial species. *Pseudomonas aeruginosa*, another Gram-negative species known for its high efflux activity, shows a similar trend: the efflux-deficient strain, in which six efflux pump systems are knocked out (Δ6) ^54^, displayed a significantly lower *k*off value compared to WT (Fig. 2c and Supplementary Fig. 7).

Together, these results demonstrate that the impact of efflux pumps on intracellular drug–target unbinding kinetics is observable at the single-cell level, largely independent of drug concentration, and conserved across bacterial species—underscoring the robustness of our findings.

A statistical mechanics model reveals that drug molecules frequently rebind to their targets in cells.

The AcrAB-TolC efflux pumps are embedded in the bacterial envelope, while the HCT’s target (DNA) resides in the cytoplasm. This spatial separation raises an important mechanistic question: how can efflux pumps influence the unbinding kinetics of drug–DNA complexes from a distance?

To explore this, we considered how the cellular environment might alter the fate of unbound drug molecules. In vitro measurements of *k*off are typically performed in dilute solutions, where drug molecules, after unbinding, diffuse away and are effectively lost to the bulk environment ^16^. In contrast, our measurements were conducted in cells, where diffusion occurs within a space bounded by the membrane. If the membrane were to behave as an absorbing boundary— removing any molecules that reach it —the fate of unbound drug molecules would resemble that of in vitro conditions.

Brownian motion of a diffusing particle in a sphere with a fully absorbing boundary is a classic problem in statistical physics. The average time a particle spends inside a sphere, the mean residence time τ, is given by ^55^:

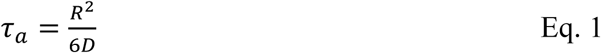

where *D* is the diffusion coefficient of the particle and *R* is the radius of a sphere. For small-molecule drugs (≲1 kDa) ^34–36^, *D* generally lies in the range of *D* = 10^0^–10^2^ µm²/s ^56,57^. For this range of *D*, assuming *R* = 1 μm for a bacterial cell, τₐ is between 0.002 and 0.2 seconds. This implies that if the cell membrane acts as a perfectly absorbing boundary, after a drug molecule unbinds from its target, it would be lost from the cell in under a second.

However, biological membranes do not behave as perfect sinks. Instead, they present a permeation barrier, reflecting a significant fraction of diffusing molecules. Whether an intracellular molecule is reflected back into the cytoplasm or crosses the membrane into the extracellular space depends on membrane permeability. At higher permeability, more molecules escape the cell and are lost. At lower permeability, more molecules are reflected back, increasing their likelihood of remaining intracellular. As a result, the residence time of a unbound drug molecule within the cell τ will depend on the permeability, although it is expected to be higher than that under an idealized absorbing boundary condition (τₐ).

To quantify this effect of the permeability on τ, we developed a statistical mechanics model of partially reflected Brownian motion (Supplementary Text, section 4). In this framework, particles diffuse within a confined compartment (e.g., a spherical bacterial cell) and encounter a boundary that probabilistically allows reflection or crossing, as defined by the Robin boundary condition (Eq. S20). Reflected particles remain within the compartment, while those that cross are irreversibly lost.

We analytically solved this model and plotted the in-cell residence time τ, normalized by τₐ, in Fig. 3a. This ratio quantifies how much longer an unbound molecule stays within a cell, relative to the fully absorbing boundary condition. This relative residence time decreases with increasing boundary permeability *κ* (Fig. 3a), since higher permeability increases the probability of escape. When *κ* exceeds approximately 10 µm/s, the ratio plateaus, indicating that the boundary behaves effectively as an absorbing surface (grey region in Fig. 3a).

**Figure 3.**
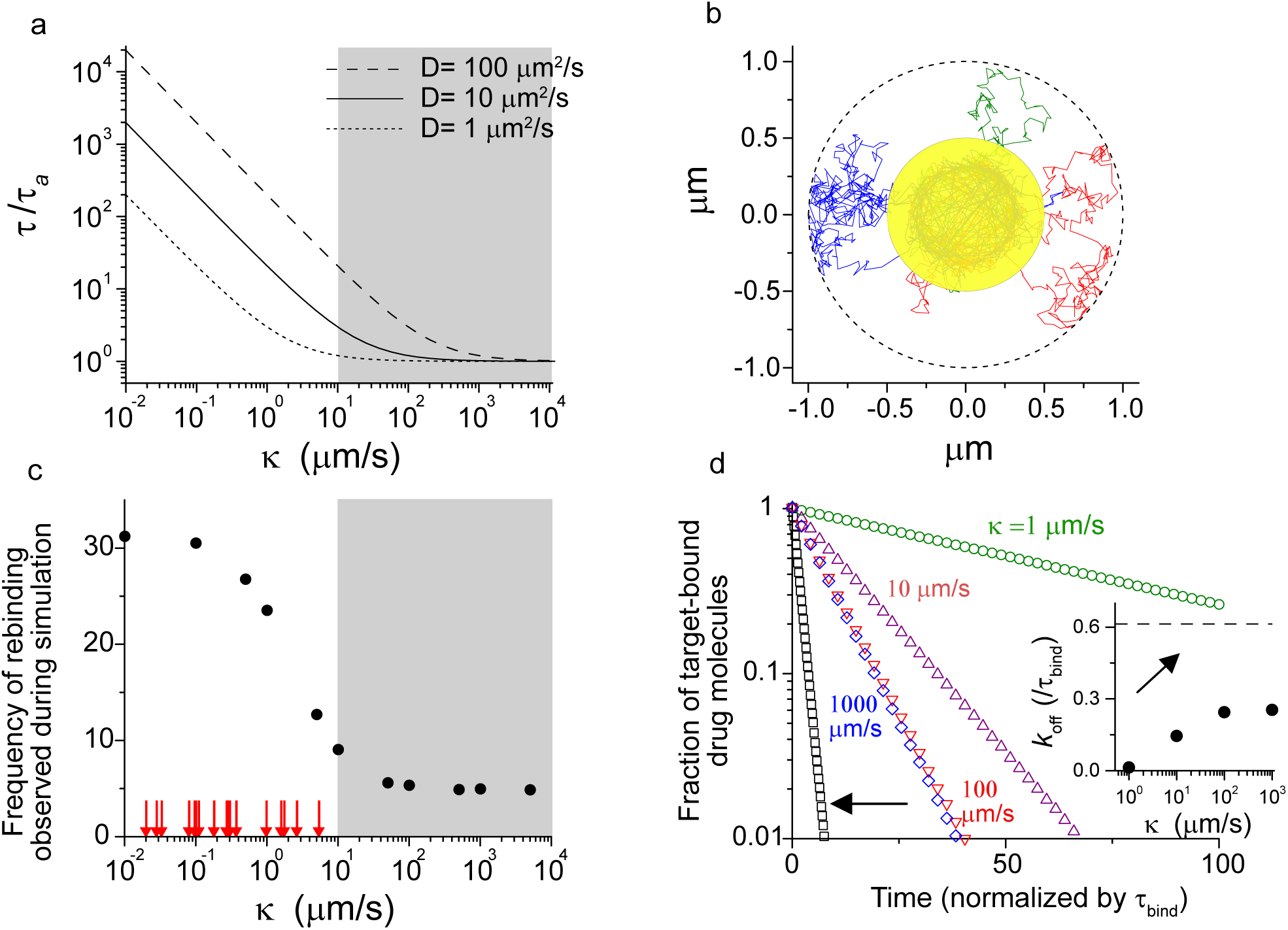
Statistical mechanics of partially reflected Brownian motion. a) The mean intracellular residence time τ of a drug molecule in a spherical cell (R=1 µm) was analytically computed as a function of boundary permeability *κ* (Eq. S30) and normalized by τa, the residence time in a cell with a fully absorbing boundary (Eq. 1 and Eq. S31). See Supplementary Text section 4 and 5 for model details. Diffusion coefficient *D* is typically in the range of 10^0^–10^2^ µm²/s ^56,57^ for small-molecule drugs (≲1 kDa) ^34–36^. The grey shade indicates the region where the boundary behaves effectively as an absorbing surface. b) A harmonic potential well at the center of a spherical cell (yellow region) represents the HCT intracellular target site, i.e., chromosomal DNA compacted into a nucleoid. Based on previous measurements ^61,62^, we set the radius of a nucleoid Rin = 0.5 µm. *D* = 10 µm^2^/s and *κ* = 1 µm/s were used for simulations. The spring constant of the harmonic well was 800/s. With this constant, the average time that a particle takes to leave the potential well, τbind, was 0.47s (Eq. S35). Typical trajectories show particles that leave the well repeatedly return to the well. c) Average number of times that a particle returns to the target site after escaping during 20 sec of simulation. 10^5^ trajectories were analyzed. Red arrows indicate experimentally measured membrane permeabilities of various drug molecules compiled from the literature (Supplementary Table). d) To simulate an unbinding experiment, particles were initialized in the central potential well representing fully bound drug–target complexes. The number of particles remaining in the well was tracked over time at various boundary permeabilities *κ*. The time is normalized by τbind. An arrow indicates a control simulation where rebinding was disabled—i.e., particles were not allowed to return to the potential well once escaped. Inset: Apparent *k*off, extracted from the slope of the decay curves. The dashed line indicates 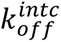 from the control simulation.

A high τ/τₐ ratio indicates that an intracellular drug molecule remains longer in the cell. A major consequence of this extended residence is an increased likelihood that unbound molecules re-encounter their targets, resulting in repeated **rebinding**. To visualize the rebinding events, we expanded our model to incorporate drug–target interactions (Supplementary Text: section 5). We introduced a harmonic potential well within a spherical cell to represent the target site (i.e., nucleoid for HCT, denoted by the yellow area in Fig. 3b). Binding was modeled as a particle becoming trapped in the well, while unbinding occurred via spontaneous exit driven by thermal fluctuations.

Example trajectories (Fig. 3b) illustrate that, after exiting the potential well, particles frequently return to the target site via diffusion and become re-trapped, demonstrating that drug molecules undergo repeated rebinding following unbinding. Because rebinding is mediated by diffusion, it occurs rapidly, on a timescale comparable to that required for a molecule to traverse the cell volume (Supplementary Fig. 8). We quantified the rebinding frequency—the number of times a unbound molecule returns to the target after initially escaping. As expected, increasing the diffusion coefficient led to more frequent rebinding (Supplementary Fig. 9a).

We next investigated the effects of boundary permeability *κ* on rebinding frequency. Above, we observed a higher τ/τₐ ratio at lower permeability, indicating that drug molecules remain longer in the cell (Fig. 3a). This extended residence time leads to more frequent rebinding events (Fig. 3c). Conversely, when *κ* increases, rebinding frequency decreases, reaching a plateau when the boundary effectively behaves as an absorbing surface (grey zone in Fig. 3c).

To assess the relevance of these findings, we surveyed published measurements of membrane permeability for various small-molecule drugs in the literature (Supplementary Table). These values, shown as red arrows in Fig. 3c, all fall below 10 µm/s—well within the regime of frequent rebinding (to the left of the grey region). This suggests that rebinding is a common feature of drug molecules in cellular environments.

In summary, these simulations highlight a kinetic competition at the core of the rebinding process; intracellular diffusion favors the return of unbound molecules to their targets, while membrane permeation promotes escape and irreversible loss. Most small-molecule drugs exhibit high diffusion coefficients (*D* = 10^0^–10^2^ µm²/s) ^56,57^ but low permeability (Supplementary Table), resulting in frequent rebinding.

### Our model predicts significant effects of rebinding on *k*off

We next investigated how rebinding influences experimentally measured *k*off (i.e., apparent *k*off). In washout experiments, *k*off is determined by first saturating the target sites with drug molecules and then monitoring a decline in target-bound molecules due to unbinding. However, frequent rebinding of unbound molecules (Fig. 3c), occurring on short timescales (Supplementary Fig. 8), can slow this decline, altering *k*off.

To quantify this effect, we computationally simulated *k*off measurements. We initialized all particles in the central potential well, representing target-bound drug molecules. We then tracked the number of particles remaining trapped in the well over time, which exhibited a first-order decay (Fig. 3d). We extracted the apparent *k*off from the slope of this decay (Fig. 3d inset). As predicted, at low permeabilities—where rebinding is frequent—the number of particles in the well declined slowly, leading to lower *k*off (Fig. 3d). Conversely, as the boundary permeability *κ* increased, which suppresses rebinding (Fig. 3c), the apparent *k*off increased (Fig. 3d).

To isolate the effect of rebinding, we ran control simulations where particles were not allowed to re-enter the potential well after escape. The resulting intrinsic unbinding rate, 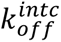, is independent of rebinding, representing the thermodynamic stability of the drug–target complex. This intrinsic rate (dashed line and arrow in Fig. 3d) was substantially higher than the apparent *k*off. Collectively, these simulation results demonstrate that rebinding significantly reduces the observed unbinding rate in cellular environments.

### Experimental tests of rebinding

The unbinding rate *k*off is traditionally regarded as an intrinsic property determined by the thermodynamic stability of drug–target complexes ^18,19^. However, our model shows that in the cellular context, when drug molecules unbind from their targets, they frequently return and rebind, causing *k*off to deviate from the intrinsic unbinding rate, 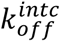. Our model also provides insight into how efflux pumps influence this deviation. Efflux pumps facilitate the export of intracellular drug molecules ^4^. This facilitated export, effectively acting as increasing the permeability for intracellular molecules, should reduce the in-cell residence time of drug molecules and hence their rebinding (Fig. 3a and 3c), increasing the apparent *k*off (Fig. 3d).

We sought to test this model prediction by quantifying 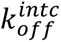 and comparing it to the apparent *k*off. Our experimental design was motivated by our simulation, where 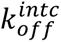 was obtained by initializing particles in the potential well and preventing their return after unbinding (Fig. 3d). In experiments analogous to this simulation, cells were first pre-loaded with HCT such that DNA was initially bound by HCT. To prevent HCT rebinding, we subsequently added an excess of Netropsin (NET), a DNA-targeting drug that occupies the same minor groove sites as HCT ^58^. As a result, the DNA sites vacated by unbinding HCT will be rapidly occupied by NET. This prevents HCT rebinding, enabling us to experimentally quantify 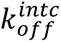.

We observed a significantly faster HCT fluorescence decay in the presence of excess NET (Supplementary Fig. 10), compared to our previous measurements in the absence of NET (Fig. 2). The slope of this decay, 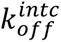, was much higher than the previously observed *k*off (compare Fig. 4a and Fig. 2b), confirming that the apparent unbinding rate deviates from its intrinsic rate in the cell. Importantly, 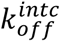 was nearly identical in WT and Δ*tolC E. coli* strains (Fig. 4a), indicating that the efflux did not alter the thermodynamic stability of the HCT–DNA complex.

**Figure 4.**
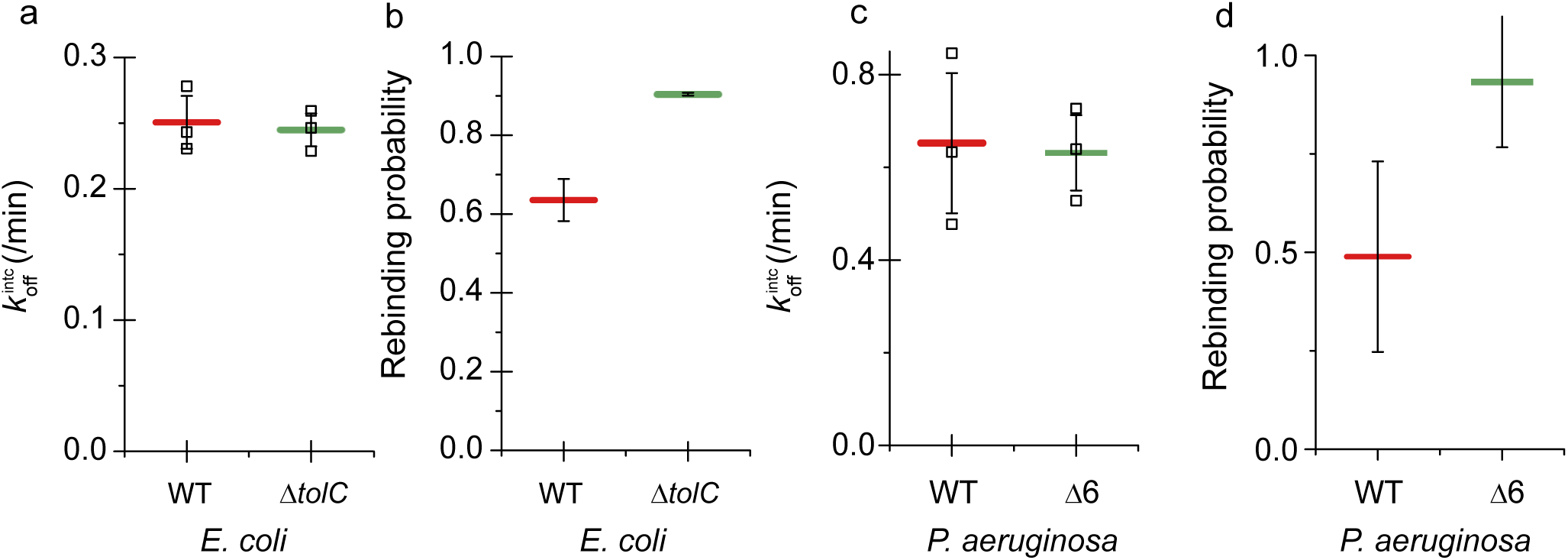
Efflux pumps modulate apparent unbinding rates by suppressing drug–target rebinding. (a) Measurement of the intrinsic unbinding rate 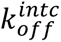 of the HCT–DNA complex. Cells were preloaded with HCT to saturate DNA sites and then exposed to excess Netropsin (NET), a competitive DNA-binding molecule, to block HCT rebinding after washout. HCT fluorescence decay was monitored over time (Supplementary Fig. 10), and 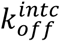 was determined from the slope of the decay. No significant difference was observed between WT and Δ*tolC* strains, confirming that efflux pumps do not alter the intrinsic chemical stability of the HCT–DNA complex. (b) Rebinding probability, deduced from the deviation of the apparent *k*off (measured without NET) from intrinsic 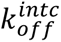 (measured with NET) using Eq. S18. The Δ*tolC* strain exhibited a significantly higher rebinding probability than WT. (c,d). 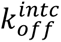 and rebinding probability in *P. aeruginosa*. Open squares represent individual values from independent replicates. Horizontal bars and error bars indicate the means and standard deviation from the replicates.

We observed the same trend in *P. aeruginosa*; HCT fluorescence decayed more rapidly in the presence of NET (compare Supplementary Fig. 11 and 7). 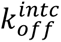 was markedly higher than the apparent *k*off (compare Fig. 2c and Fig. 4c). In particular, 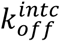 was similar between WT and Δ6 strains (Fig. 4c).

We next determined the degree of rebinding in efflux-active and deficient cells. We have observed in our computer simulation that rebinding causes the deviation of apparent *k*off from 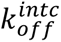 (Fig. 3d). We mathematically formulated the relationship between this deviation and rebinding probability (Eq. S18). Then, inserting the measured values of *k_off_* and 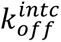 (Fig. 2bc and 4ac) into this equation, we quantified the rebinding probability. The probability was lower in efflux-active strains than in their efflux-deficient counterparts, in both *E. coli* and *P. aeruginosa* (Fig. 4b and 4d), confirming our model prediction that efflux pumps suppress rebinding.

### Differences in *k*off result in different drug–target affinities

Another often-cited parameter of drug efficacy is the drug–target binding affinity, quantified as an equilibrium dissociation constant *K*D 16. *K*D is equal to the ratio of the unbinding and binding rates, *K*D =*k*off / *k*on. Therefore, any change in *k*off should directly influence the drug-target affinity inside the cell. In particular, a higher *k*off in efflux-active strains relative to efflux-deficient strains (Fig. 2) should lead to a higher *K*D, indicating weaker binding affinity.

While *K*D could be deduced from the kinetic constants (*k*off and *k*on), it can also be quantified independently of them by measuring the ligand-receptor complex concentrations at the steady state against ligand concentrations, where the *K*D equals 50% of the maximum response ^16^. We measured the HCT–DNA *K*D through this independent approach. Intracellular HCT fluorescence intensity indicates the concentration of DNA-bound HCT molecules. We measured the steady state intensity *F*s across a range of external HCT concentrations (Supplementary Fig. 12). The dose–response curve was well explained by a standard binding model, which we then fit to extract the apparent *K*D (Supplementary Fig. 12 and caption).

We found that the apparent *K*D was approximately 4.5-fold lower in Δ*tolC* cells compared to WT (Supplementary Fig. 12), closely matching the ∼4-fold difference in *k*off between these two strains (Fig. 2b). Given that *K*D =*k*off / *k*on, this match suggests that the increased binding affinity in Δ*tolC* arises primarily from the reduction in unbinding rate, rather than changes in the association rate. Collectively, our data indicate that efflux pumps affect the drug-target equilibrium constant (affinity) by altering the unbinding kinetics, mediated through a change in rebinding frequency.

## Discussion

Drug–target interactions, characterized by their binding affinity and kinetic rates, are central to determining drug efficacy ^16,19^. In our study, we observed a lower drug-target affinity and higher unbinding rates, i.e., weaker drug-target interactions, in efflux-active cells than efflux-deficient cells. Following the common explanation in in vitro studies, we had initially considered whether this effect was due to altered thermodynamic stability of drug–target complexes. However, our direct measurements of the intrinsic unbinding rate, 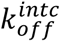, confirmed that the thermodynamics of their interactions remain unchanged (Fig. 4).

To resolve this paradox, we conducted in-depth live-cell measurements and statistical-mechanics-based modeling. The results revealed that drug–target interactions in the cellular environments are stabilized by rapid rebinding of drug molecules. Efflux pumps limit rebinding by removing unbound drug molecules from the cell, thereby destabilizing drug-target interactions through a kinetic—not thermodynamic—mechanism.

This finding uncovers a previously unrecognized mode of efflux-mediated resistance. For a drug to be intracellularly functional, it must not only enter the cell but also occupy its targets. Any impediments to either 1) drug entry or 2) target occupation contribute to drug resistance ^2^. Efflux pumps are well known for their “gatekeeping” role in limiting drug entry. Consistent with this, we observed that they reduce intracellular HCT accumulation by approximately fivefold (Fig. 1 and Eq. S12). The focus of this study was on the latter, demonstrating that efflux pumps also reduce target occupation remotely. In particular, this “housekeeping” mechanism increased the HCT-DNA unbinding rate by approximately 4-fold and the apparent affinity by a similar degree (Fig. 2 and Supplementary Fig. 12).

This 5-fold restriction in HCT entry and 4-fold reduction in affinity together explains why a 20-fold (=5×4) higher external HCT concentration was required in WT cells to achieve the same level of target occupancy to that of Δ*tolC* (Fig. 1a–c). Thus, efflux pumps do not merely block drug entry—they actively reshape intracellular drug-target interactions. Fully understanding their impact requires accounting for synergistic interactions between these two functions, underscoring the multifaceted and dynamic nature of efflux-mediated drug resistance.

Our quantitative analysis provides an important mechanistic insight into this housekeeping function. Most drugs are small molecules (≲1 kDa) ^34–36^, with the diffusion constant generally in the range of *D* = 10^0^–10^2^ µm²/s ^56,57^. This rapid diffusion leads to frequent rebinding to their targets (Fig. 3). Our model further examined the interplay between this intracellular diffusion and boundary permeability, identifying the regime where the frequency of rebinding is high but sensitively dependent on the permeability *κ* (Fig. 3c). To put this finding in a perspective, we compiled experimentally measured membrane permeabilities of various small-molecule drugs (Supplementary Table, red arrows in Fig. 3c). Remarkably, most of their values fall within the above identified regime of high frequency/sensitivity. This overlap indicates that efflux pumps are well poised to limit drug-target rebinding in the cell across a broad spectrum of drug molecules. Coupled with their broad substrate specificity—particularly those of the RND family ^8^—this finding provides new physical insights into how efflux pumps contribute broadly to multi-drug resistance.

These insights also have important implications for understanding drug action and designing new therapeutics. While extensive efforts have focused on optimizing the thermodynamic properties of drug–target interactions ^18,20,59,60^, the accessible chemical space for such optimization is not infinite. This work underscores the importance of spatial constraints and kinetic factors in intact cells—such as cellular dimensions, intracellular diffusion, membrane permeability, and efflux activity—in shaping drug efficacy. By altering these factors, it may be possible to enhance drug performance without modifying their chemical properties, offering a novel strategy for improving therapeutic outcomes.

## Data Availability Statements

Source data for figures are provided in the Source Data file.

## Code Availability Statements

Our custom-built software is provided as Supplementary file.

## Acknowledgements

This work was funded by NIH (1U19AI158080, SD, KS, DL, MK). We thank Graeme Conn for sharing the *Pseudomonas aeruginosa* strain deficient in efflux pumps (PAO1Δ6).

## Author Contributions

MK conceived the study. SD, KS and DL designed and carried out the experiments. SD developed the mathematical model. MK secured funding and provided resources. MK and SD wrote the manuscript. All authors read and approved the manuscript.

## Methods

### Bacterial strains and culture conditions

*E. coli* K-12 NCM3722 was used ^63,64^. Δ*tolC* was constructed using P1 transduction using Keio collection knockouts ^65^. *Pseudomonas aeruginosa* strain PAO1 and PAO1Δ6 ^54^ (Δ*mexAB-oprM,* Δ*mexCD-oprJ* Δ*mexEF-oprN* Δ*mexJKL,* Δ*mexXY,* Δ*triABC*) were gifts from Graeme Conn lab (Emory University).

Cells were cultured in Neidharts MOPS minimal media ^66^ with 10 mM glucose and 10 mM NH4Cl as the carbon and nitrogen sources.

To prepare experimental cultures, cells were taken from −80°C stocks and streaked on a LB plate. A single colony was inoculated in 2 mL LB medium and grown at 37°C with constant agitation at 250rpm in a water bath. To monitor growth, the optical density (OD600) of the culture was measured using a Genesys20 spectrophotometer with a standard quartz cuvette. Before cells entered stationary phase, cells were inoculated into 5 mL MOPS minimal medium at very low densities (typically lower than the OD600 of ∼0.0001) and cultured overnight (pre-culture). Next morning, the pre-culture was diluted in pre-warmed, 3 mL MOPS minimal medium (experimental culture) to the OD600 of ∼0.01 and allowed to grow exponentially.

### Hoechst 33342 measurements

Hoechst 33342 (HCT) was purchased from Sigma Aldrich. HCT was added to the exponentially growing culture at OD 0.05 at desired concentrations. Samples were taken at specified time points and placed on a microscope coverslip. A 1.5% agar pad containing the same medium composition as the culture was placed on the coverslip and transferred to the microscope stage.

Images were acquired using an Olympus inverted fluorescence microscope with a 60× oil lens. Fluorescence was taken with a DAPI filter set using 500 msec exposure time. Brightfield images for identification of whole cells were captured at the same time using 200 msec exposure through the same filter cube and phase contrast illumination. At least 30 cells were imaged. Images were recorded using an Andor Neo 5.5 CMOS camera and analyzed using imageJ ^67^. Thresholding was conducted on the phase contrast images to identify cells, creating a mask. This mask was then overlayed on the fluorescence images, and the average fluorescence was calculated.

In experiments to measure *k*off, 1 mM and 0.05 mM HCT for *E. coli* (WT and Δ*tolC*) and 50 µM and 15 µM for *P. aeruginosa* (PAO1 and PAO1Δ6) was added to culture and incubated for at least 30 min. The culture was then centrifuged at 2000g for 5 min, the supernatant replaced with fresh medium lacking HCT, centrifuged again to wash out any residual HCT. Samples were taken at specific time points and measured as described above.

To measure 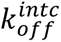, a growing culture was inoculated with HCT at OD600 of 0.03 for 30 minutes. 5 µg/ml Netropsin (Enzo Life Sciences) for *E.coli* and PAO1, 2.5 µg/ml for *E. coli* Δ*tolC*, 0.3 mg/ml for PAO1Δ6 was added and incubated for 20 mins. HCT was removed from the medium by centrifugation and replaced with Netropsin-containing media.

### Simulation

We performed a Monte Carlo simulation in C++. The particle position was updated iteratively at each time step using Eq. S33 within a spherical harmonic potential well and Eq. S34 outside the potential well. The simulation space was confined within a spherical boundary of radius *R*. When the particle reached this boundary, whether it was absorbed or reflected was determined probabilistically using Eq. S20, with the probability of absorption given by 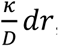, where κ is the permeability coefficient and *dr* is the radial increment. If absorbed, the simulation for that particle was terminated; otherwise, the particle was reflected and repositioned within the boundary. The time step Δ*t* was selected to satisfy the stability condition 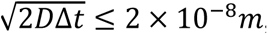, ensuring that the particle’s displacement remained within the smallest spatial resolution used, thereby maintaining numerical accuracy throughout the simulation. The force constant *k* was set to 800 *s*^−1^ throughout the simulation. This value was selected because it provides a balance where the average residence time of the particle within the harmonic potential well is neither too short nor too long. To ensure statistically robust estimates of key metrics such as the residence probability within the well, the return time distribution, and the frequency of rebinding events, we simulated 10^5^ independent particle trajectories. For a comprehensive description of the simulation framework and parameter choices. See Section 4 and 5 in the Supplementary Text for details.

