## Supplementary Material for "Beyond Gatekeeping: Efflux Pumps Remotely Destabilize Cytoplasmic Drug–Target Interactions by Limiting Rebinding"

**Supplementary Text**

**Section 1: Modeling the permeation barrier for drug influx.**

Although drug permeation in the presence of efflux pumps was quantitatively modeled in several studies [1-3], these models contained numerous parameters, which makes it difficult to understand the effects of efflux pumps. Here, we adopt a coarse-grained approach.

We first consider the bi-directional membrane permeation of drug molecules without active transports (e.g., efflux-deficient strains). Let $P$ denote the effective permeability of drug molecules. For an external drug concentration, $C_{out}$, the total influx rate of drug molecules, $J_{in}$, is given by

$$J_{in} = P \times C_{out}. \qquad \text{Eq. S1}$$

Likewise, for an internal drug concentration $C_{in}$, total outflux rate of drug molecules, $J_{out}$, is

$$J_{out} = P \times C_{in}. \qquad \text{Eq. S2}$$

A change in the intracellular drug concentration is governed by their balance,

$$\frac{dC_{in}(t)}{dt} = P(C_{out} - C_{in}(t)), \qquad \text{Eq. S3}$$

Immediately after drug molecules are introduced to the media, $C_{in}$ is low. Thus, the equation above can be approximated as

$$\frac{dC_{in}(t)}{dt} = PC_{out}, \qquad \text{Eq. S4}$$

or

$$C_{in}(t) = PC_{out} \times t. \qquad \text{Eq. S5}$$

Therefore, immediately after drug addition, the initial rise in $C_{in}(t)$ can be approximated as a linear function, as we have shown in Fig. 1a and 1b. The slope of this rise is equal to $PC_{out}$.

After this initial rise, $C_{in}$ will ultimately reach a steady state. Setting $\frac{dC_{in}(t)}{dt} = 0$ in Eq. S3, we have

$$C_{in}(\infty) = C_{out}. \qquad \text{Eq. S6}$$

Efflux pumps are known to serve as a permeation barrier for drug influx [4]. We modeled this process by a reduced permeability for incoming drug molecules, $P_{inward}$.

Then, Eq. S3-S6 should be re-written as

$$\frac{dC_{in}(t)}{dt} = P_{inward} \times C_{out} - P \times C_{in}(t), \qquad \text{Eq. S7}$$

$$\frac{dC_{in}(t)}{dt} = P_{inward} \times C_{out}, \qquad \text{Eq. S8}$$

$$C_{in}(t) = P_{inward} C_{out} \times t. \qquad \text{Eq. S9}$$

$$C_{in}(\infty) = \frac{P_{inward}}{P} C_{out}. \qquad \text{Eq. S10}$$

In the main text, we showed that WT exhibits a slope that is 3.7-fold higher than Δ*tolC* (Fig. 1d).

Then, combining Eq. S5 and S10, we have

$$\frac{P \times C_{out}^{\Delta tolC}}{P_{inward} \times C_{out}^{\mathrm{WT}}} = \frac{1}{3.7}, \qquad \text{Eq. S11}$$

where $C_{out}^{WT} = 1\ \mu M$ and $C_{out}^{\Delta tolC} = 0.05\ \mu M$ in our experiments (Fig. 1). Therefore,

$$\frac{P}{P_{inward}} = \frac{1}{3.7} \times \frac{C_{out}^{\mathrm{WT}}}{C_{out}^{\Delta tolC}} = \frac{20}{3.7} = 5.4. \qquad \text{Eq. S12}$$

This indicates that the effective membrane permeability for incoming drug molecules for WT is 5.4-fold lower than that of Δ*tolC*, supporting the role of efflux pump as a permeation barrier.

**Section 2: Drug-target binding/unbinding kinetics.**

Consider free drug molecules at concentration *C*, binding reversibly to their intracellular targets. Let $T_T$ denote the total concentration of available target sites, and $T$(t) the concentration of the drug–target complex at time *t*. The kinetics of complex formation can be described by the following ordinary differential equation:

$$\frac{dT(t)}{dt} = k_{on} C (T_T - T(t)) - k_{off} T(t) \qquad \text{Eq. S13}$$

Here, $k_{on}$ and $k_{off}$ are the binding and unbinding rate constants, respectively.

In the regime where target saturation is negligible (i.e., $T(t) << T_T$), the available target pool remains effectively constant, and the equation simplifies to:

$$\frac{dT}{dt} = k_{on} C T_T - k_{off} T. \qquad \text{Eq. S14}$$

The solution is

$$T(t) = \frac{k_{on}}{k_{off}} C T_T [1 - \exp(-k_{off} t)]. \qquad \text{Eq. S15}$$

This solution shows that the time at which the steady state is reached is governed by the unbinding rate $k_{off}$.

**Section 3: Deriving the rebinding rate from unbinding kinetics.**

Consider $N$ number of HCT molecules initially bound to their target. After wash-out, $\Delta N$ molecules unbind over $\Delta t$,

$$\Delta N = -k_{off}^{intc} N \Delta t. \quad \text{Eq. S16}$$

If these unbound molecules rebind with a probability of $R$, $\Delta N = -k_{off}^{intc} N \Delta t + R \cdot k_{off}^{intc} N \Delta t$, or alternatively,

$$\frac{dN}{dt} = -(1-R) \cdot k_{off}^{intc} N. \quad \text{Eq. S17}$$

Therefore, the apparent unbinding rate $k_{off}$ differs from the intrinsic unbinding rate $k_{off}^{intc}$ by a factor of $1-R$. Alternatively, if we quantify $k_{off}$ and $k_{off}^{intc}$ (which we have shown in the main text with and without Netropsin), we can determine *R* as follows:

$$R = 1 - k_{off}/k_{off}^{intc}. \quad \text{Eq. S18}$$

**Section 4: Mean residence time of a particle inside a sphere**

Consider an intracellular drug molecule diffusing in the cytoplasm. When it encounters the cell membrane, it can either reflect back to the cytoplasm or cross the membrane, escaping the cell. We model this process by a particle undergoing Brownian diffusion with a diffusion coefficient *D* within a sphere with a radius *R*. The boundary of the sphere is partially reflective, as defined by the permeability $\kappa$. We calculated the mean residence time of a particle in a sphere, τ, i.e., how long molecules stay in the sphere on average [5].

Denoting the particle's radial position from the center by $r$, its diffusion can be modeled by a diffusion equation

$$\frac{\partial c(r,t)}{\partial t} = D \frac{\partial^2 c(r,t)}{\partial r^2} + \frac{2}{r} \frac{\partial c(r,t)}{\partial r}. \quad \text{Eq. S19}$$

$c(r,t)$ is the probability distribution function of a particle at time *t*. We assumed that this particle was initially located at the center of the sphere: $c(r,0) = \frac{\delta(r)}{4\pi r^2}$. The sphere boundary is semi-permeable and has a permeability $\kappa$, yielding a boundary condition:

$$-D \frac{\partial c}{\partial r}\Big|_R = \kappa c(R,t). \quad \text{Eq. S20}$$

To solve the equation, we eliminate the singularity at the origin by defining a new function $u(r,t) = rc(r,t)$, which reduces Eq S19 to

$$\frac{\partial u}{\partial t} = D\frac{\partial^2 u}{\partial r^2}, \quad \text{Eq. S21}$$

and Eq. 20 to

$$-D\frac{\partial u}{\partial r}|_{r=R} = \kappa u(R,t). \quad \text{Eq. S22}$$

Applying separation of variables with $u(r,t) = R_n(t)T_n(t)$, we obtain solutions of the form $T_n(t) = e^{-Dk_n^2 t}$ and $R_n(r) = sin\,(k_n\, r)$, where the constants $k_n$ are determined by applying the boundary condition.

Substituting the spatial eigenfunctions $u_n$ into the above equation leads to a transcendental equation for the eigenvalues $k_n$

$$k_n R cot k_n R = 1 - \frac{\kappa R}{D}. \quad \text{Eq. S23}$$

Thus, the general solution for the probability distribution function takes the form

$$c(r,t) = \sum_1^\infty A_n \frac{sink_n r}{r} e^{-Dk_n^2 t}. \quad \text{Eq. S24}$$

To determine the coefficients $A_n$, we make use of the orthogonality of the spatial eigenfunctions $\sin k_n R$ over the interval [0, R], weighted appropriately for the radial problem. Applying Eq. S24 to the initial condition, we get

$$c(r,0) = \frac{\delta(r)}{4\pi r^2} = \sum_1^\infty A_n \frac{sink_n r}{r}. \quad \text{Eq. S25}$$

We multiply both sides by $\frac{4\pi r^2 \sin k_m r}{r}$ and integrating from 0 to $R$ and using the condition $\delta_{mn} = 0$ (when $m \neq n$), we get

$$A_n = \frac{k_n}{4\pi F(k_n R)}, \quad \text{Eq. S26}$$

where $F(k_nR) = \int_0^R \sin^2 k_n r\, dr$. Substituting these coefficients back into Eq S25, we obtain the full time-dependent probability distribution function:

$$c(r,t) = \sum_1^\infty \frac{\mathrm{k_n}}{4\pi F(k_nR)} \frac{sin k_n r}{r} e^{-Dk_n^2 t}. \quad \text{Eq. S27}$$

where the eigenvalue $k_n$ is the solution of Eq. S23.

The survival probability, $S(t)$, which quantifies the probability that a particle remains within the spherical region at time $t$, is obtained by integrating the probability density over the volume of the sphere:

$$S(t) = \int_0^R 4\pi r^2 c(r,t) dr. \quad \text{Eq. S28}$$

The mean residence time, τ, is defined as the expected time a particle spends inside the sphere before being absorbed. The term, $-\frac{dS(t)}{dt}$ refers to the fraction of a particle absorbed between *t* and *t+dt*, i.e., the probability distribution of a particle residing in the sphere up to time *t*. Therefore,

$$\tau = -\int_0^\infty t \frac{dS(t)}{dt}\, dt = \int_0^\infty S(t) dt. \quad \text{Eq. S29}$$

Substituting the series expression for $c(r,t)$ in Eq S27 into the above definition and evaluating the integrals term by term yields a compact expression for the mean residence time:

$$\tau = \frac{2\kappa R^3}{D^2} \sum_1^\infty \frac{sin k_n R}{k_n^3 R^3 \left(1 - \frac{\sin 2k_n R}{2k_n R}\right)}. \quad \text{Eq. S30}$$

When $\kappa$ is infinite, the boundary behaves like an absorbing boundary. Eq S30 reduces to the mean residence time for a complete absorbing boundary, $\tau_a$

$$\tau_a = \frac{R^2}{6D}. \quad \text{Eq. S31}$$

**Section 5: Simulating particle dynamics in a harmonic trap inside a sphere**

We next expanded the above model to include drug-target binding/unbinding to the DNA complex in the cytoplasm. Binding of a drug molecule to its target is modeled by a trap in harmonic potential well.

In the model, a three-dimensional harmonic potential well with a radius $R_{in}$ is placed inside the sphere (Fig. 3b): $R_{in} < R$. This potential well is concentric with the sphere. A particle in the potential well experiences a restoring force of strength $k$.

The motion of the particle inside the potential well is described by the following stochastic differential equation:

$$d\vec{X} = -k\vec{X}dt + Dd\overrightarrow{W(t)}. \quad \text{Eq. S32}$$

The first term corresponds to the deterministic restoring force directed towards the center of the potential well. The second term describes the effects of thermal fluctuations, where $D$ is the diffusion coefficient and $\overrightarrow{W(t)}$ represents the Wiener process increments [6], capturing the stochastic nature of the particle's trajectory.

Unbinding of a drug molecule from its target is captured by the escape of the particle from the harmonic potential well.

Once the particle escapes this well, its motion is described by the model in S.4), where a particle's motion is governed by pure diffusion, with the outer boundary permeability $\kappa$.

To study the diffusion dynamics of a particle, we performed a Monte Carlo simulation using C++. The simulation proceeds by initializing the particle's position at the origin ($X = 0$). At each time step, the particle's position is updated based on the above stochastic equation. Inside the harmonic potential well, the particle's displacement is calculated separately for each spatial direction, namely the $X$, $Y$and $Z$ coordinates. The displacement equation in each direction ($x_i$, where $i = 1, 2, 3$) is given by

$$x_i(t + \Delta t) = x_i(t) + kx_i dt + \sqrt{2D\Delta t}\xi(t). \quad \text{Eq. S33}$$

Here, $x_1$, $x_2$ and $x_3$ correspond to the particle's positions along the $X$, $Y$and $Z$ axes, respectively. $\xi(t)$ represents a random number drawn from a normal distribution with mean zero and variance one. Outside the harmonic well, the position update follows the equation

$$x_i(t + \Delta t) = x_i(t) + \sqrt{2D\Delta t}\xi(t). \quad \text{Eq. S34}$$

If the particle reaches the spherical boundary at $r = \sqrt{x_1^2 + x_2^2 + x_3^2} = R$, a probabilistic decision based on the permeability $\kappa$ determines whether the particle is absorbed or reflected. Following the boundary condition at Eq S20, the probability of absorption is given by $\frac{\kappa}{D}dr$, where $dr$ is the increment in radial direction. If reflected, the position is adjusted to remain within the boundary; if absorbed, the simulation for that particular particle ends.

The time step for the simulation, $\Delta t$, is chosen based on the diffusion coefficient $D$ used in each specific simulation. To ensure numerical stability and accuracy, the condition $\sqrt{2Ddt} \leq 2 \times 10^{-8}m$ is imposed, where $2 \times 10^{-8}m$ represents the smallest spatial step selected for our simulation. This criterion ensures that the displacement per time step remains within physically reasonable limits, preventing numerical errors.

To calculate the probability that a particle resides inside the potential well, we performed simulations of multiple particle trajectories. Specifically, we generated over $10^5$ independent trajectories to ensure statistical accuracy. At each time point, we computed the residence probability by determining the fraction of particles that remain within the harmonic well. This approach allows us to capture the time-dependent decay of the residence probability with high precision.

The average time that a particle remains trapped within the harmonic potential well, denoted as $\tau_{bind}$. can be calculated analytically. This quantity represents the average binding time between a Hoechst molecule and DNA, reflecting the stability of the Hoechst-DNA complex. To obtain $\tau_{bind}$, we consider the situation where the boundary at $R_{in}$ is a completely absorbing boundary. In this configuration, the particle follows the stochastic equation given by Eq. S32. Starting from the origin, the mean first passage time to the absorbing boundary at $R_{in}$ corresponds to the average time it takes for the particle to escape the potential well, i.e., $\tau_{bind}$. For an Ornstein–Uhlenbeck process in three dimensions as in our model in Eq S32, this mean first passage time can be calculated using a well-established theoretical framework [7]:

$$\tau_{bind} = \frac{1}{D}\int_0^{R_{in}} dz \frac{e^{\frac{k}{2D}z^2}}{z^2} \int_0^z t^{-2} e^{-\frac{kt^2}{2D}} dt \qquad \text{Eq. S35}$$

This analytical expression accounts for the combined effects of the harmonic potential's restoring force and the thermal diffusion of the particle within the trap.

We also measured the mean return frequency of the particle to the DNA complex after it exits the harmonic potential well. To quantify this, we tracked the number of times the particle returned to the potential well after escaping, for each individual trajectory. The simulation was conducted over a large number of independent trajectories ($10^5$) to ensure statistical robustness. The mean return frequency was then calculated by averaging the number of returns across all trajectories. For each simulation, we monitored the particle's movement for up to 20 seconds to capture multiple return events.

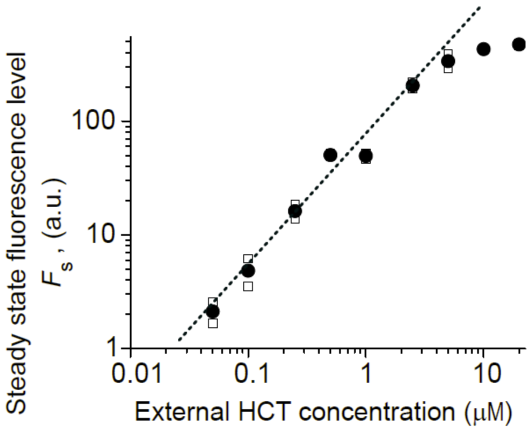


**Supplementary Fig. 1. Steady-state fluorescence level, $F_s$, at different external HCT concentrations.**

As shown in Fig. 1a, upon HCT addition (time zero), the fluorescence intensity in the cells, *F*(t), initially increased and reached a steady-state level around 15 mins. We repeated this experiment using various HCT concentrations. The data points between 15 and 30 mins were averaged to obtain the steady-state fluorescence level, $F_S$. Open squares represent individual values from two independent replicates. Solid symbols represent their means. Above 200 a.u., increasing external HCT concentration only minimally increase $F_S$, indicating that it is in the saturating regime.

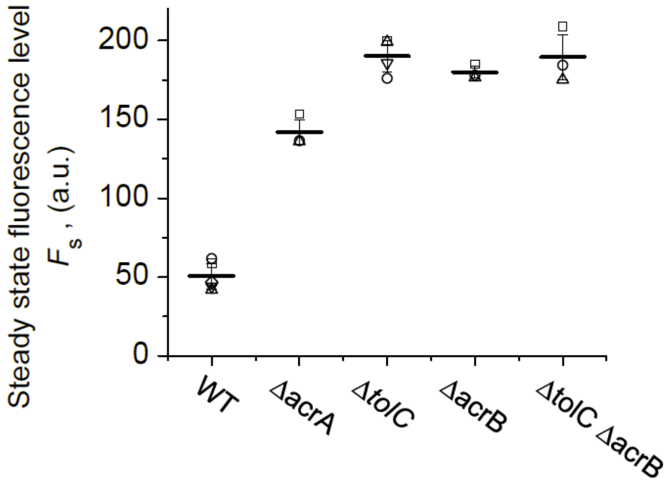


**Supplementary Fig. 2. Steady state HCT intensity, $F_s$, in WT, Δ*acrA*, Δ*acrB* and Δ*tolC*.**

The AcrAB-TolC system is composed of a tripartite complex including the outer-membrane channel (TolC), the inner membrane transporter (AcrB), and the periplasmic adaptor protein (AcrA) [8]. A single knockout of *acrA* increased $F_s$, although to a lesser extent than single knockouts of *acrB* or *tolC*. The double knockout strain (Δ*tolC* Δ*acrB*) exhibited an $F_s$ comparable to that of the corresponding single knockouts. Fluorescence intensity was measured 30 minutes after HCT addition. A concentration of 1 μM HCT was used for all strains. Open squares represent individual values from three independent replicates. Horizontal bars represent their means. Error bars indicate their standard deviations. At 1 μM HCT, Δ*tolC* and Δ*acrB* showed fluorescence levels near 200 a.u., indicating they are in the saturation regime (see Supplementary Fig. 1).

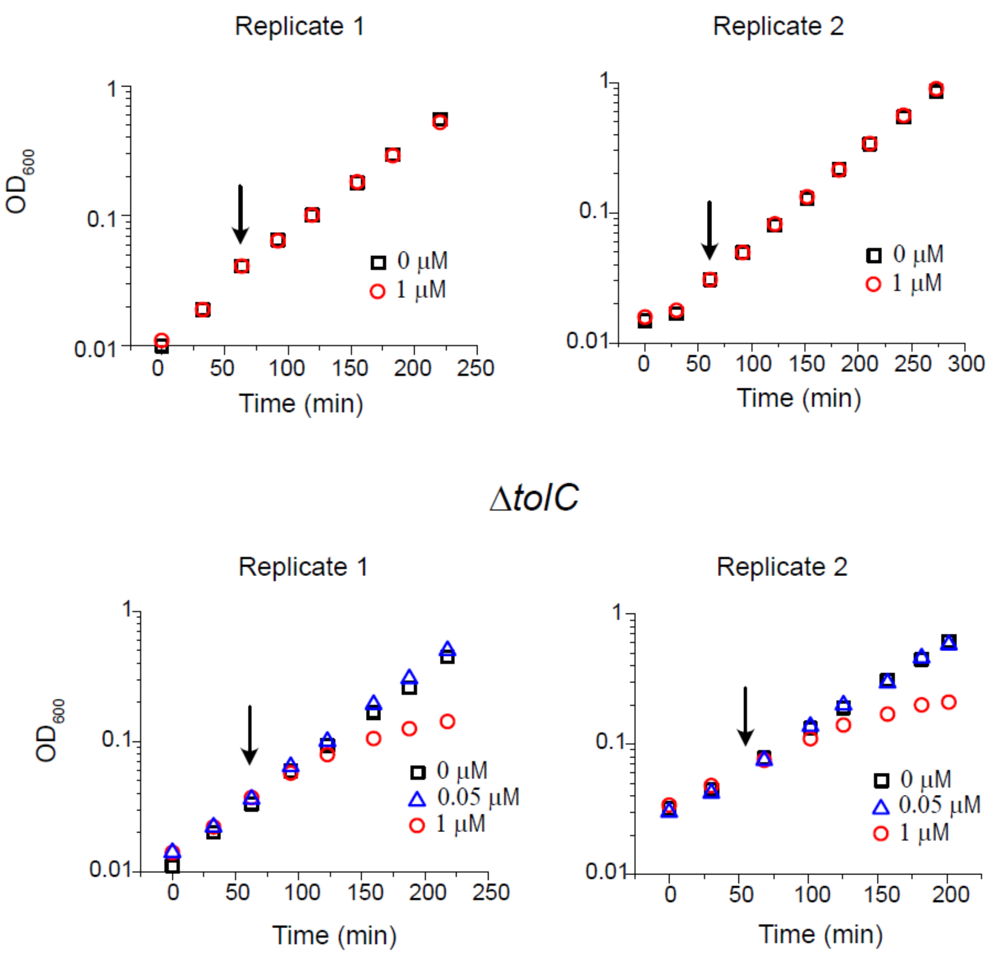


**Supplementary Fig. 3. WT and Δ*tolC* growth rates at different HCT concentrations.**

1 µM HCT did not affect WT growth whereas it significantly reduced the Δ*tolC* growth. The Δ*tolC* growth was not affected at 0.05 µM HCT.

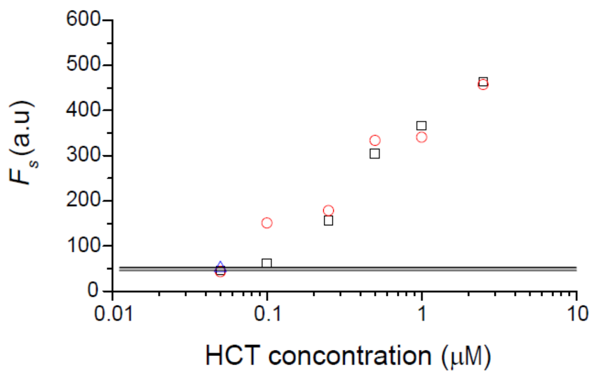


**Supplementary Fig. 4. Steady-state fluorescence intensity ($F_s$) in the Δ*tolC* strain across a range of external HCT concentrations.**

$F$(t) were measured at 20, 25, and 30 minutes after HCT addition and averaged to estimate $F_s$. The horizontal line indicates the $F_s$ level observed in WT cells at 1 μM HCT for comparison.

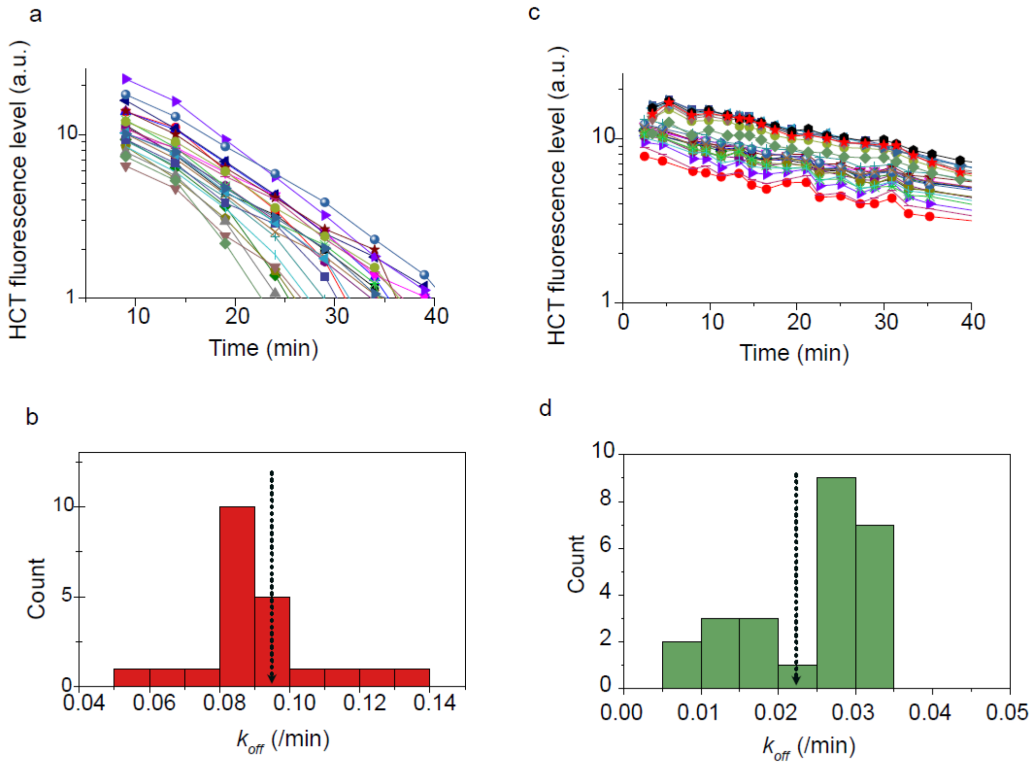


**Supplementary Fig. 5. Single-cell measurements of HCT–DNA unbinding kinetics.**

Time-lapse microscopy was used to track intracellular fluorescence decay in individual WT (a,b) and Δ*tolC* cells (c,d) following HCT washout. The unbinding rate constant $k_{off}$ was determined for each cell. The arrows indicate the population-level $k_{off}$ values (Fig. 2b), confirming that the observed difference in unbinding kinetics is not due to population heterogeneity.

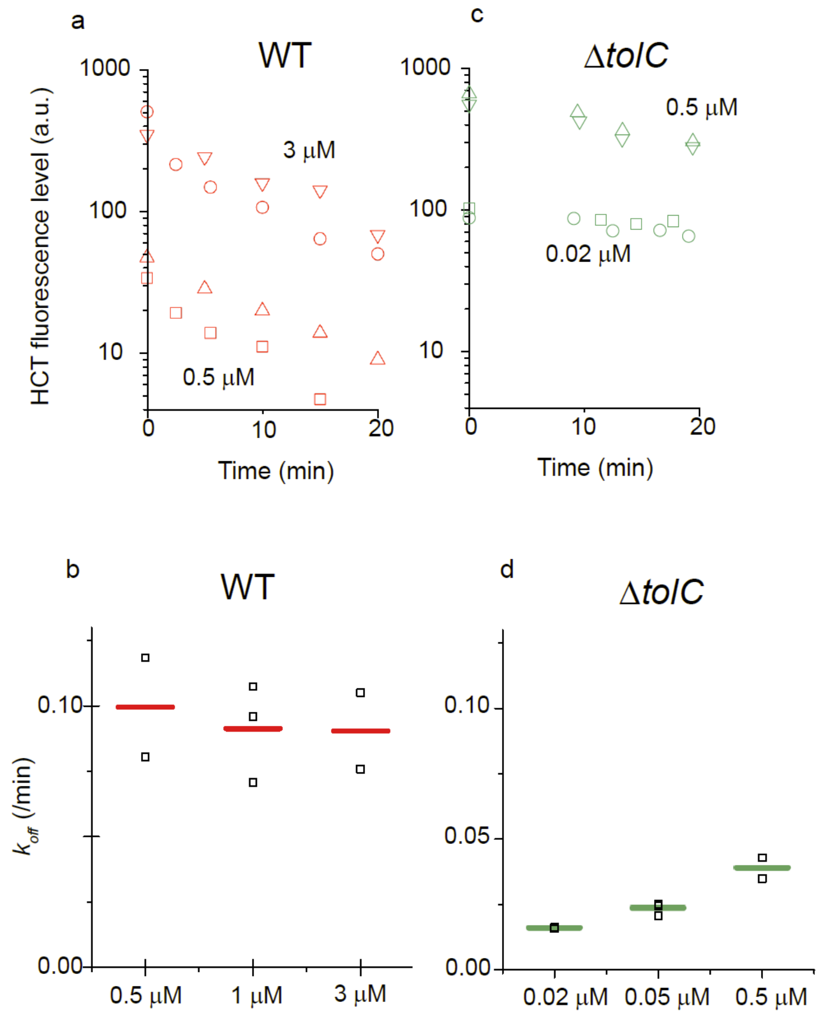


**Supplementary Fig. 6. Dependence of HCT–DNA unbinding rate on external HCT concentration.**

The unbinding rate $k_{\text{off}}$ was measured in WT and Δ*tolC* strains across a range of external HCT concentrations. Only minor variation in $k_{\text{off}}$ was observed. Δ*tolC* consistently exhibited a significantly lower unbinding rate than WT. Two or three biological replicates were conducted (different symbols). Horizontal bars indicate the means.

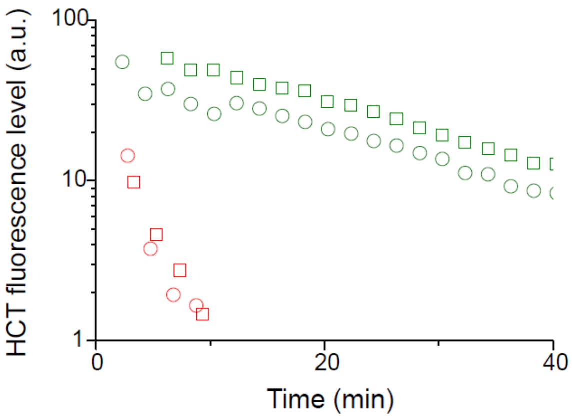


**Supplementary Fig. 7.** HCT fluorescence level upon washout in *P. aeruginosa*

After allowing HCT fluorescence to reach its steady-state level, we washed the cells and suspended them in HCT-free media. HCT fluorescence decay was monitored over time. The experiments were repeated twice (symbols with different shapes). Red/green symbols indicate efflux-active and efflux-deficient strains respectively. See Fig. 2c and Methods for the experiment and strain details.

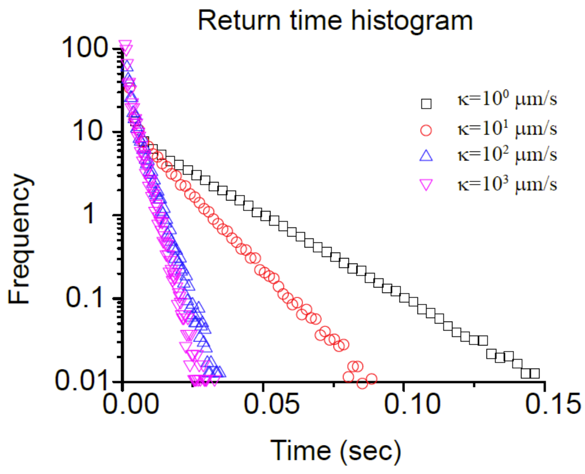


**Supplementary Fig. 8. Return time histogram for various boundary permeabilities *κ*.**

In the main text, we estimated that a freely diffusing small molecule can traverse the bacterial cell volume in a fraction of a second; $\tau_a = 0.02$ seconds for $D = 10^{-11}$ m²/s. We analyzed the particles trajectories (as exemplified in Fig. 3b) to determine how quickly particles that once left the central well return to the well. The plot shows that most of the rebinding occurs in the estimated diffusion timescale.

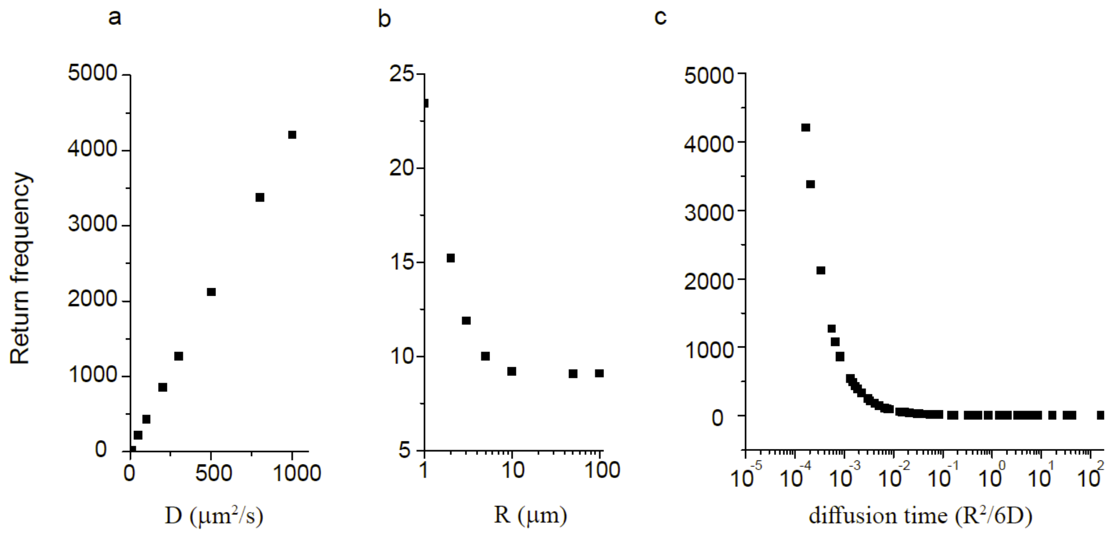


**Supplementary Fig. 9. Return frequency as a function of *D*, R, and diffusion time.**

Repeating our simulation, we calculated on average how frequently a molecule returns to the central potential wall in 20 second long trajectories for different *D* and R values. The boundary permeability remained fixed ($\kappa$ = 10 μm/s). a) *D* was varied (R = 1μm). b) R was varied (*D* = 10 μm$^2$/s). c) The time it takes for a molecule to diffuse spherical volume is given by $\frac{R^2}{6D}$ (Eq. 1). We collapsed our data by this diffusion time.

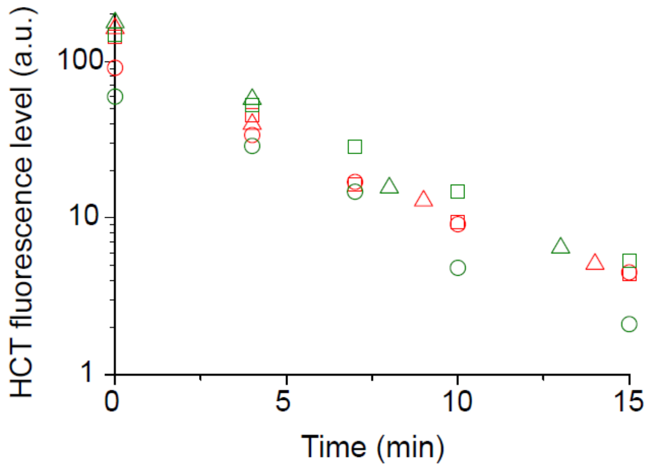


**Supplementary Fig. 10.** HCT fluorescence level upon wash-out in the presence of Netropsin.

Cells were preloaded with HCT and then exposed to an excess of Netropsin (NET), a competitive DNA-binding molecule, to block HCT rebinding after washout. HCT fluorescence decay was monitored over time. The experiments were repeated three times (symbols with different shapes). Red/green symbols indicate WT/Δ*tolC*, respectively.

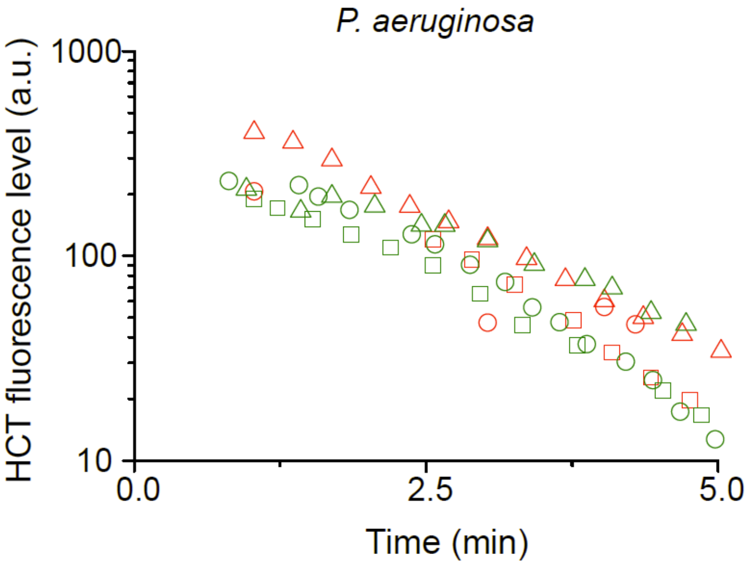


**Supplementary Fig. 11.** HCT fluorescence level upon wash-out in the presence of Netropsin in *P. aeruginosa*.

The experiments were conducted and data plotted as described in Supplementary Fig. 10.

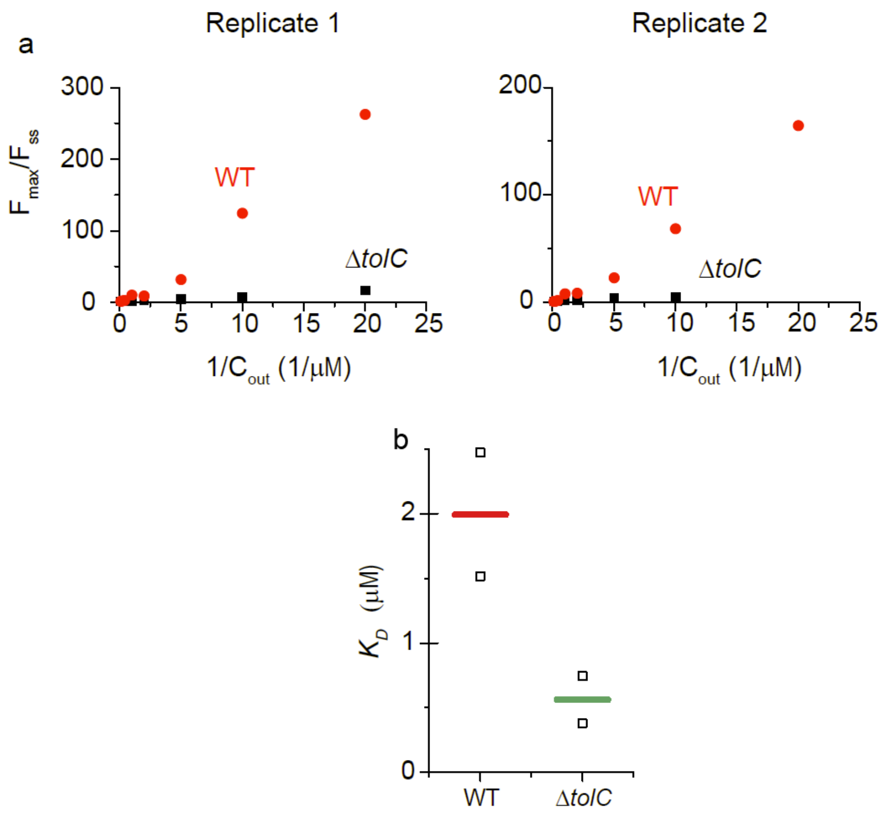


$K_D$ in μM

| | WT | Δ*tolC* |
|---|---|---|
| Replicate 1 | 2.5 | 0.74 |
| Replicate 2 | 1.5 | 0.38 |
| Mean | 2.0 | 0.56 |

**Supplementary Fig. 12.** Calculating HCT-DNA binding affinity $K_D$ for WT and Δ*tolC* strains.

We measured the steady state fluorescence intensity, $F_s$ at different external HCT concentrations, $C_{out}$. To determine $K_D$ from this relationship, we consider target occupancy, i.e., the number of DNA minor grove target sites occupied by HCT at the steady state, $T_S$, divided by the total number of target sites, $T_T$. Setting $\frac{dT(t)}{dt} = 0$ in Eq. S13, together with $K_D = \frac{k_{off}}{k_{on}}$ [9], we have

$$\frac{T_S}{T_T} = \frac{C_{in}}{K_D + C_{in}}, \quad \text{Eq. S36}$$

HCT molecules fluoresce only when they bind to DNA [10-13]. Assuming that their fluorescence level is proportional to the number of target-bound HCT, we have

$$\frac{T_S}{T_T} = \frac{F_S}{F_{max}}, \qquad \text{Eq. S37}$$

where $F_{max}$ is the maximum fluorescence level we observed at high $C_{out}$ (10 µM); see Supplementary Fig. 1.

We therefore have

$$\frac{F_{max}}{F_S} = 1 + K_D \frac{1}{C_{in}}. \qquad \text{Eq. S38}$$

Describing this equation in terms of $C_{out}$ using Eq. S6 and S10, we have

$$\frac{F_{max}}{F_S} = 1 + K_D \frac{1}{C_{out}} \qquad \text{for } \Delta tolC \qquad \text{Eq. S39}$$

and

$$\frac{F_{max}}{F_S} = 1 + K_D \frac{P}{P_{inwward}} \frac{1}{C_{out}} \qquad \text{for WT} \qquad \text{Eq. S40}$$

where $\frac{P}{P_{inwward}} = 5.4$ from Eq. S12.

It predicts a linear relationship between $\frac{F_{max}}{F_S}$ and $\frac{1}{C_{out}}$, which agrees with our observations (The panel a). Importantly, the slope informs the $K_D$ (panel b and the table below).

**Supplementary Table:** Membrane permeability for various antibiotics.

| Antibiotic | MW (g/mol) | P (µm/s) | Reference |
|---|---|---|---|
| Imipenem | 299 | 1.75 | [14] |
| Benzylpenicillin | 333 | 0.007 | [15] |
| Penicillin G | 334 | 0.02 | [16] |
| Ampicillin | 349 | 0.028, 0.98 | [15,16] |
| Cefaclor | 368 | 2.63 | [16] |
| Carbenicillin | 378 | 0.0075 | [16] |
| Meropenem | 383 | 0.3 | [14] |
| Cephaloram | 389 | 0.08 | [17] |
| Cephalothin | 395 | 0.29, 0.11 | [16,17] |
| Cephaloglycine | 406 | 0.98 | [17] |
| Cephaloridine | 415 | 5.37, 3.57 | [16,17] |
| Cefoxitin | 427 | 0.37 | [16] |
| Azthreonam | 435 | 0.033 | [16] |
| Cefazolin | 453 | 1.58, 0.62 | [16-18] |
| Cefotaxime | 455 | 0.18 | [16] |
| Cefamandole | 461 | 0.1 | [17] |
| Cephamandole | 462 | 0.11 | [16] |
| Piperacillin | 516 | 0.27 | [14] |
| Cefsoludin | 531 | 0.18 | [17] |
| Cefsulodin | 533 | 0.3 | [16] |
| Ceftazidime | 546 | 0.096 | [16] |
| Erythtomycin | 734 | 0.00021 | [3] |
| Tetracycline | 444 | 0.000056 | [3] |
